# Ultrasensitive quantification of HIV-1 cell-to-cell transmission in primary human CD4^+^ T cells measures viral sensitivity to broadly neutralizing antibodies

**DOI:** 10.1101/2023.09.08.556871

**Authors:** Dmitriy Mazurov, Alon Herschhorn

**Affiliations:** Division of Infectious Diseases and International Medicine, Department of Medicine, University of Minnesota, Minneapolis, MN 55455, USA; Institute for Molecular Virology; Institute for Engineering in Medicine; Center for Genome Engineering, University of Minnesota, Minneapolis, MN 55455, USA; Microbiology, Immunology, and Cancer Biology Graduate Program; The College of Veterinary Medicine Graduate Program; Molecular Pharmacology and Therapeutics Graduate Program, University of Minnesota, Minneapolis, Minnesota 55455, USA

**Author notes:** Correspondence: Alon Herschhorn Room 2-101 MRF 689 23^184^ Av SE, University of Minnesota Minneapolis, MN 55455.

## Abstract

HIV-1 efficiently replicates *in vivo* by direct transmission from infected to uninfected CD4^+^ T cells at confined local sites designated virological synapses (VSs). VSs are formed by cell junctions between HIV-1 envelope glycoproteins (Envs) on an infected cell and CD4 on an uninfected cell. These sites facilitate highly efficient viral transmission and contribute to HIV-1 evasion from neutralizing antibodies, but accurate quantification of the efficiency of cell-cell transmission is still challenging. Here, we developed an ultrasensitive HIV-1 cell-to-cell transmission assay that triggers the expression of the *nanoluciferase (nluc)* gene in target cells upon transmission and after reverse transcription of the HIV-1 RNA genome. The assay is based on insertion of the *nluc* gene in an antisense orientation into HIV-1 provirus; Nluc expression is blocked in virus-producing cells because the *nluc* gene contains an intron that can be efficiently spliced out only when mRNA is transcribed from the opposite (sense) strand. Thus, only sense mRNA that is spliced and subsequently reversed transcribed during transmission to target cells will support Nluc expression. Assay optimization resulted in a very low background, >99% splicing efficiency, high sensitivity and wide dynamic range for detection of cell-cell transmission in T cell lines and primary CD4^+^ T cells. The new reporter vector can detect cell-cell transmission using single-round viral vectors and HIV-1 molecular clones, which provide viral proteins of different HIV-1 strains, and reproducibly measures sensitivity of HIV-1 transmission to antibody neutralization *in vitro*. This assay will contribute to understanding fundamental mechanisms of HIV-1 cell-to-cell transmission, allow evaluation of pre-existing or acquired HIV-1 resistance in clinical trials, and can be adapted to study other retroviruses.

## INTRODUCTION

HIV-1 replicates in host cells to produce new progeny that will then infect new target cells by one of two alternatives modes: 1) budding virions diffuse into biological fluids, where they are diluted, and circulate in the organism until they encounter new permissive cells, and 2) an infected cell contacts directly a permissive cell and form virological synapses (VSs), where the budding virions are concentrated in close proximity to the membrane of an uninfected cell and can efficiently interact with the CD4 and CCR5/CXCR4 receptors to enter the new target cell (1). Mechanisms of HIV-1 cell-cell transmission typically involve changes in producer cells, including engagement of adhesion molecules on infected cells with related molecules on uninfected target cells and cytoskeleton reorientation towards cell-cell contact. However, despite wealth of information, aspects of HIV-1 cell-cell transmission through the VSs such as signaling and activation of target cells, membrane sites of the virion budding and fusion at VSs, and correlation between morphological events and HIV-1 infection are still controversial or poorly understood.

Accurate measurements of HIV-1 cell-cell transmission in vitro is challenging. Immunofluorescent staining of HIV-1 Gag transfer to target cells using microscopy or flow cytometry, which is often used to quantify the cell-cell transmission of HIV-1 in primary cells (2), does not necessarily reflect productively infectious events (3). The use of fluorescent molecular clones such as iGFP (4) to monitor cell-cell transmission requires additional labeling of target cells. And conventional luciferase-based single- or replication competent HIV-1 assays express high levels of reporter protein in transfected cells, which are used to produce the viruses for transmission, that obscure the detection of newly infected target cells in co-culture assays typically used for cell-cell transmission experiments *in vitro*. To overcome some of these limitations, we have previously developed the replication-dependent reporter vectors that confine expression of the reporter protein to post transmission stage and after reverse transcription of HIV-1 genome in the newly infected cell (Figure 1A). These vectors commonly contain an expression cassette consisting of a promoter, an intron-containing reporter gene, and polyA signal; the complete cassette is inserted in the antisense orientation relative to the viral sequence (5). This specific design prevents the expression of active reporter protein in transfected (virus-producing) cells from the two mRNAs that are transcribed (Fig. 1A): expression from LTR-mediated mRNA transcript is blocked because the reporter gene is positioned in a reverse orientation, and expression from the internal promoter (CMV)-mediated mRNA is blocked because the mRNA contains the complement splicing sites and therefore the intron cannot be efficiently spliced out. Only when the viral genome, which is transcribed from the LTR, is efficiently spliced, packaged into the nascent virion and reverse transcribed during transmission to target cell the reporter protein is expressed (Figure 1A). One drawback of this system is the dependency on splicing of the reporter gene in the viral genome since both spliced and unspliced mRNA can be packaged into the nascent virions, and the latter does not support reporter protein expression (Figure 1B). Subsequent efforts optimized the intron in the reporter gene and used anti-intron shRNA to reduce unspliced mRNA packaging, creating in-GFPturbo and in-mCherry vectors that resulted in up to 80% spliced mRNA packaging in HIV-1 virions (6). However, the splicing efficiency for mRNA from similar vectors containing the *firefly luciferase* gene remained low and did not allow accurate and sensitive measurements of HIV-1 cell-cell transmission in human primary CD4+ T cells.

**Figure 1.**
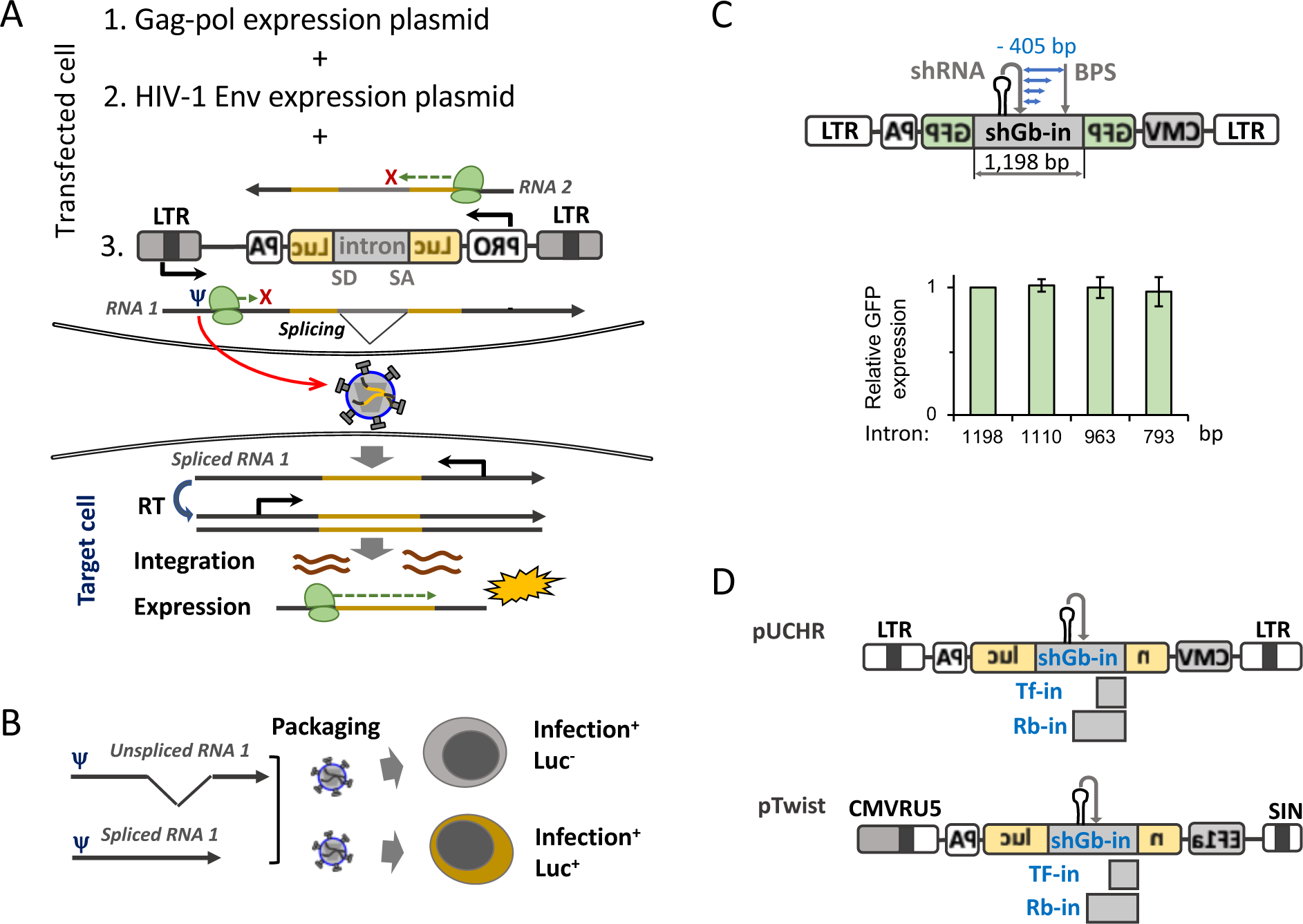
Design of cell-cell transmission reporter vectors. (**A**) A scheme of engineered elements in the reporter vector. These include the following blocks in the reverse orientation: promoter (PRO), 5’- and 3-portions of reporter *luc* gene separated by an intron, and polyA (PA) signal (all shown by inverted letters). The splice-donor (SD) and the splice-acceptor (SA) sites flanking the intron are not reversed and therefore mRNA transcribed from the promoter cannot be spliced. Transfection leads to transcription of 2 mRNAs: RNA 1 can be spliced but will not express the reporter protein because the reading frame is reversed; RNA 2 cannot be spliced and therefore will not express the reporter protein. Only RNA 1 can be packaged into virions due to the ψ signal and during productive infection, reverse transcriptase (RT) will synthesize (-) strand DNA containing the *luc* gene in the correct orientation, (+) strand DNA and integrates the double stranded DNA into host chromosome where the Luc protein can be expressed from the promoter PRO. (**B**). One potential limitation of using the reporter vector is the packaging of unspliced RNA 1 into the nascent virions. This will decrease detection cell-to-cell transmission since the resulting integrated provirus will still contain the intron and will not support Luc expression. (**C**) Top, a scheme of shortening of shRNA-modified human γ-globin intron (shGb-in) between shRNA target site and branch point site (BPS). Bottom, the effect of shortening shGb-in on cell-cell transmission readout in 293T cells using vesicular stomatitis virus G (VSV_G) protein to mediate viral entry. (**D**) A scheme of new nanoluciferase (*nluc)*-based lentiviral vectors with three types of introns generated and tested in this study: shGB-in, mouse TNFβ intron (TF-in), and ribozyme intron (Rb-in). Luc – firefly luciferase; LTR – long terminal repeat; CMV-cytomegalovirus promoter; EF1a – elongation factor 1a promoter, SIN – self-inactivating 3-LTR. Results shown in panel C are the mean ± SD of three independent experiments.

Importantly, HIV-1 cell-cell transmission exhibits varied degree of resistance to broadly neutralizing antibodies (bnAbs) against HIV-1. These bnAbs target highly conserved regions of the envelope glycoproteins (Envs) and block viral entry of different HIV-1 strains (7–12). bnAbs are developed in a small fraction of people living with HIV-1 after several years of infection and some bnAbs prefer to neutralize specific Env conformations (13–15). Immunotherapy and antibody-mediated protection trials are currently evaluating bnAbs as treatment of prevention modalities, but the documented ability of HIV-1 cell-cell transmission to evade bnAbs *in vitro* is a significant concern for the application of bnAbs to medical interventions. Thus, a robust and reproducible assay to monitor cell-cell transmission in primary CD4+ T cells could provide new tools to understand the mechanisms of HIV-1 transmission between cells and to investigate mechanisms of HIV-1 resistance to bnAbs by cell-to-cell transmission.

Here, we designed and built a reversed intron reporter vector (designated in-Nluc) to measure HIV-1 cell-cell transmission that is based on the *nano luciferase* gene and depends on HIV-1 for replication. Splicing efficiency of mRNA transcribed from in-Nluc is higher than 99% and sensitivity is >10^4^-fold higher than the original firefly luciferase vectors. in-Nluc reporter quantitatively measures HIV-1 cell-to-cell transmission in different cell-coculture systems (e.g., from adherent 293T to Cf2-Th/CD4^+^CCR5^+^ cells, and between T cell lines), including peripheral blood CD4^+^ T cells. High dynamic range and low background enables accurate measurements of inhibition of HIV-1 cell-cell transmission by neutralizing antibodies. in-Nluc will be a helpful tool both for the evaluation of HIV-1 resistance in clinical trials and for fundamental studies of the mechanisms of HIV-1 cell-to-cell transmission.

## MATERIAL AND METHODS

### Cell cultures, primary cell isolation and activation

Human embryonic kidney 293T cells were obtained from the American Tissue Culture Collection (ATCC), Cf2-Th/CD4^+^CCR5^+^ cells stably expressing human CD4 and CCR5 were kindly provided by Joseph Sodroski (Dana-Farber Cancer Institute), CEM cells were obtains through NIH AIDS Reagent Program, and SupT1.CCR5 cells stably expressing the human CCR5 coreceptor were a kind gift from James Hoxie (University of Pennsylvania). Trima Cones containing concentrated human leukocytes from freshly collected blood were purchased from Innovative Blood Resources (Minneapolis, USA). Trima cone cells were diluted 1:3 with PBS, and the peripheral blood mononuclear cells (PBMCs) were isolated on the density gradient of lymphocyte separation medium 1077 (PromoCell, Heidelberg, Germany) using SepMate 50 ml tubes (StemCell Technologies, USA) and then directly used or frozen in fetal bovine serum (FBS; Gibco) containing 5% DMSO (Sigma, USA). Before using in downstream applications, frozen PBMCs were thawed and incubated in a culture medium for one day. CD4^+^ T cells were isolated from fresh or frozen PBMCs by negative selection using the EasySep™ human CD4+ T cell isolation kit according to the manufacturer’s instructions (Stem Cell Technology, USA), and activated with CD2/CD3/CD28 magnetic beads (Miltenyi Biotec, Germany) for 48 hr. Immediately prior to electroporation, the activating beads were removed from CD4^+^ lymphoblasts using the magnet. 293T and Cf2-Th/CD4^+^CCR5^+^ cells were cultured in Dulbecco’s modified Eagle’s medium (DMEM) (Sigma-Aldrich, USA) with 10% FBS, 2 mM glutamine, 100 U/ml penicillin, and 100 µg/ml streptomycin (all from Gibco, ThermoFisher). CEM, SupT1.CCR5 cells were maintained in RPMI 1640 medium containing 10% FBS, 2 mM glutamine, 100 U/ml penicillin, and 100 µg/ml streptomycin. Activated human CD4^+^ T cells were cultured in RPMI-1640 growth medium supplemented with 50 IU/ml of recombinant human interleukin-2 (Miltenyi Biotech, Germany). Cells were maintained at low passage and tested periodically for mycoplasma contamination.

### Plasmid construction

We optimized the length of the intron of the human γ-globin gene by generating 3 short PCR-amplified fragments using separately 3 forward primers, each combined with the common reverse primer, as indicated in Supplementary Table S1, and pUCHR-inGFPt-mR plasmid (6) containing the full-length intron as a template. The resulting three PCR fragments of 104, 251, and 421 bps were separately cloned back into the pUCHR-inGFPt-mR plasmid via the XbaI/SpeI restriction sites. The nano luciferase-based HIV-1 reporter vectors pUCHR-inNluc and pTwist-inNluc containing shortened ψ-globin intron with shRNA (shGb), intron from the mouse TNFβ gene (Tf), and *Tetrahymena* ribozyme intron (Rb) were designed as described in the Results section by Gene Universal, Delaware. The sequences of expression cassettes for each reporter vector are provided in Supplementary Figure 1. To generate the pUCHR-EF1a-inNluc plasmid we subcloned the hybrid EF1a-HTLV promoter from the pTwist-Tf-inNluc into pUCHR-inNluc plasmid, replacing the CMV promoter. The pTwist-Tf-inNluc plasmid was digested with Sal I, blunted, and then digested with Nhe I; the resulting digested insert was subcloned into the SmaI/NheI sites of pUCHR-inNluc. Plasmids containing the molecular clones HIV-1_NL4-3_ (pNL4-3), HIV-1_NL4-AD8_ (pNL(AD8)), and the packaging plasmid psPAX2 were obtained from the NIH AIDS Reagent Program; the packaging vector pCMV-dR8.2 was from Addgene. Plasmid containing HIV-1_NL4ΔEnv_ (pNL4ΔEnv) sequence, which carry a frame-shift mutation due to re-ligation of Nde I bunted ends in *env* gene, was generated by digesting NL4-3 Gag-iGFP ΔEnv plasmid (NIH AIDS Reagent Program) with Sal I / Xho I and cloning the resulting DNA fragment into the same restriction sites of pNL4-3 plasmid. The Env expression plasmid pcDNA 3.1-AD8-M, pSVIIIe7-AD8, pcDNA 3.1-KB9 and pCMV-VSVG (expression vector for protein G of vesicular stomatitis virus) were from Joseph Sodroski (Dana-Farber Cancer Institute). The firefly luciferase expression plasmid pGL3-CMV was purchased from Promega (Madison, WI).

### Quantification of spliced and unspliced reporter RNA packaging

To determine the splicing efficiency, 293T cells (5×10^5^ cells per well of a 6-well plate) were cotransfected with 0.35 μg of psPAX2 or pNL(AD8) and 0.45 μg of a reporter plasmid using Effectene transfection reagent (Qiagen, USA). The next day, culture medium was removed, cells were gently washed with warm media once, and incubated in 1.5ml of the medium for another day. Supernatants were collected, centrifuged at 3,000 x g for 5 min to remove cell debris, and further centrifuged (in 1.5 ml tubes) at 18,000 x g for 2.5 h to isolate viral particles. Viral RNA was isolated from the particles using QIAamp DSP Viral RNA Mini Kit (Qiagen, USA), and treated with RNAse-free DNAse I (New England Biolabs) to remove traces of the plasmid DNA. All purified viral RNA was reverse transcribed using random hexamer and SuperScript III reverse transcriptase (Invitrogen, USA) for 1 h at 50°C after a short 10-min preincubation at 25°C. cDNA amount was assessed by quantitative PCR (qPCR) using GoTaq qPCR Master Mix (Promega, USA), 0.4 μM of each primer, and 1-5 ng of cDNA template. The primers used for amplification of unspliced reporter RNAs (intron), spliced reporter RNAs, and HIV-1 genomic RNA (conserved Gag region) are shown in Supplementary Table S2. PCR was performed using CFX-96 Real-Time PCR Detection System (Bio-Rad, USA) at the following settings: one cycle of denaturation at 95°C for 2 min and 40 cycles of amplification (95°C 7 sec, 58.4°C 15 sec, and 60°C 15 sec). Data were collected and analyzed by using CFX-96 software. Levels of spliced and unspliced reporter RNAs in viral particles were quantified for each transfer vector. The efficiency of RNA splicing was expressed as a percentage of the ratio of spliced to the total RNA (spliced and unspliced) and was calculated by the formula:

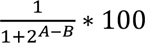

, where A is a threshold cycle (*C_T_*) for spliced RNA, and B is *C_T_* for unspliced RNA (6). When molecular clones were used, a similar calculation was performed to determine the percentage of spliced reporter RNA relative to the total (spliced, unspliced reporter, and viral genomic) RNAs packaged into HIV-1 virions

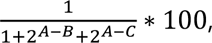

, percentage of unspliced reporter RNA relative to total three RNAs:

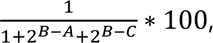

, or percentage of HIV-1 genomic RNA relative to total three RNAs:

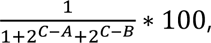

where C is *C_T_* of HIV-1 genomic RNA.

### Cell-cell transmission assay with coculturing of 293T and Cf2-Th/CD4^+^CCR5^+^ cells

HEK 293T cells or HEK 293T cells mixed with Cf2-Th/CD4^+^CCR5^+^ cells at 1:1 ratio were added to wells of a white, flat bottom 96-well plate (Greiner, USA) in 100 μl of complete DMEM at a concentration of 2×10^5^ cells/mL. The plate was incubated overnight and then cells were cotransfected with 10 ng reporter plasmid, 8 ng psPAX2, and 2 ng HIV-1 Env- or VSV-G-expression plasmid. pcDNA3.1 was used as a negative control instead of Env expression plasmids. In some experiments, the reporter plasmid was cotransfected with 8 ng pNL(AD8). Cells were transfected in triplicates using the Effectene transfection reagent according to manufacturer’s instructions (Qiagen). The transfection mixture was adjusted to 50 μl per well using a growth medium before adding to cells. In some experiments, the non-nucleoside reverse transcriptase inhibitor Etravirine (NIH AIDS Reagent Program) was added directly to the transfection mixture. At indicated time points or after 48-h, the supernatants were removed and 1X 50 μl of Glo Lysis Buffer (Promega, USA) were added to each well and incubated for at least 10 min. Saturated-signal samples were further diluted in Glo Lysis buffer. To measure nano luciferase activity, NanoGlo luciferase substrate (Promega, USA) was diluted 1:50 in NanoGlo Buffer, and 30 μl of the reagent was added to 50 μl of lysates. Plates were incubated at room temperature for 3 min before luciferase activity was measured with 3-sec integration time using Centro XS^3^ LB 960 luminometer (Berthold Technologies, USA).

### Cell coculture infectivity assays with T cell lines and primary lymphocytes

10^6^ CEM or activated CD4^+^ T cells were washed once with PBS (in 1.5 ml tubes), and electroporated in buffer R with 3 μg of the reporter plasmid, and 2 μg of one of the packaging plasmids psPAX2, pCMV-dR8.2 or pNL4ΔEnv, in the presence or absence of 1 μg of Env-expression plasmid. In some experiments, the reporter plasmid was cotransfected with 2 μg pNL4-3 or pNL(AD8). Cells were electroporated using Neon Transfection System (Invitrogen, USA), 100 μl tips, and either 1230V, 40 ms x 1 pulse for CEM cells, or 1600V, 10 ms x 3 pulses for CD4+ T cells. Electroporated cells were washed once in 1 ml of culture medium to remove plasmid DNA and then mixed with 5×10^5^ SupT1.CCR5 cells; CD4^+^ T lymphocytes were mixed with 5×10^5^ autologous CD4^+^ T lymphocytes in final volume of 1.5 ml growth medium in a 12-well plate for 72 hr. Fifty U/ml recombinant human interleukin-2 (IL-2) was added to cultures of primary CD4^+^ T cells. To measure Nluc activity, cells were centrifuged (in 1.5 mL tubes) at 1,000 x g for 3 min and lysed in 50 μl of Glo lysis buffer. Cell lysates were transferred to white 96-well plate, and luciferase activity was measured using NanoGlo reagent as described above for adherent cells.

### Cell coculture neutralization assay with T cell lines in a 96-well plate

Antibody was diluted in RPMI culture medium by 3-fold serial dilutions as previously described (16); control wells were medium without antibody. Transfected CEM and target SupT1.CCR5 cells, which was prepared as described above, were mixed thoroughly and 50 μl were added into the 96-well plate with diluted antibody. Plates were incubated at 37°C for 72 hr, and cells were lysed by adding 30 μl NanoGlo reagent and aspirating/dispensing several times. Luciferase activity was measured within 3-10 min after the addition of NanoGlo reagent.

### Free virus infectivity and neutralization assay with T cell lines

To quantify HIV-1 infection of viral particles produced by T cells, 0.75 ml of virion-containing supernatant obtained after centrifugation of co-cultures (12-well plates) used to assay cell-cell transmission was added to 5×10^5^ SupT1.CCR5 target cells that were resuspended in 0.75 ml fresh medium. Three days later, cells were lysed, and the level of HIV infection was measured by Nluc activity. To avoid transferring cells along with supernatants when using 96-well plates, we transfected 10^6^ CEM cells (1 ml) and grow the cells in a 12-well plate for 72 hr. Transfected cells were resuspended and centrifuged at 1,000 x g for 3 min (in 1.5 ml tubes). Two-thirds of the supernatant was harvested from the top of the sample and used for the viral neutralization assay. After 72-h incubation, levels of Nluc activity were measured as described above.

### Estimation of HIV-1 replication in reporter-transfected CD4 T cells

10^6^ CEM or CD4^+^ T cells were transfected with 5 μg of pUCHR-EF1a-inNluc reporter vector as described above, washed once with medium and incubated with 300 ng (p24) HIV-1_NL4-3_ in final volume of 100 μl for 4 hr at 37°C. After centrifugation step and removal of viral supernatants, cells were washed with 0.5 ml of warm medium, and incubated with the indicated target cells in 1.5 ml of culture medium. The medium was supplemented with IL-2 for infection of primary CD4^+^ T cells. Cell cocultures were incubated in a 12-well plate for the indicated times. Every 2-3 days, cells were suspended and 1 ml of the sample was removed for analysis and replaced with fresh medium. Cell pellets were lysed in 50 μl of Glo lysis buffer and Nluc activity was measured as described above; supernatants were used to quantify the level of HIV-1 p24 antigen by ELISA (XpressBio, Frederick, MD).

### Statistical analysis and data presentation

Data are the average + SD of at least three independent experiments. Student’s *t* test was used to calculate statistical difference between groups. Dose-response curves of viral neutralization by bnAbs were fitted to the four-parameter logistic equation using Prism 7 program (GraphPad, San Diego, CA) after adding the equation to the program; IC_50_ values and the associated s.e. are reported (17–19). The number of experiment repeats and replicates are provided in Figure Legends.

## RESULTS

### Design and development of cell-cell transmission vectors expressing nanoluciferase

Current lentiviral-based reporter vectors to detect cell-cell transmission contain HIV-1 Rev response element (RRE) for viral RNA nuclear export and a packaging signal (Ψ) for packaging the reporter RNA into virions. These requirements result in packaging of RNA that contains either spliced or unspliced reporter gene into virions, decreasing the sensitivity of the assay that requires spliced RNA to express intact reporter protein in target cells (Figure 1B). One approach to significantly increase packaging of RNA with spliced reporter gene into virions is to engineer anti-intron Mir30-shRNA *cis* element within the reporter gene intron to facilitate degradation of RNA containing unspliced reporter gene in the cytoplasm (6). However, applying this strategy to reporter vectors containing the *firefly luciferase* gene (in-Luc-mR), resulted so far in low splicing efficiency.

We designed and built an improved lentiviral-based reporter vector to efficiently and accurately measure HIV-1 cell-to-cell transmission by introducing the following modifications to current vectors (Fig. 1). **1)** we replaced the firefly luciferase gene (*fluc*) with the nanoluciferase gen (*nluc*), increasing signal readout by 1-2 orders of magnitude (20,21). **2)** we optimized the intron by testing three different introns within the *nluc* gene: a) short-length version (793 bp; Fig. 1C) of shRNA-modified human γ-globin intron (shGb-in) (6); b) intron II from the mouse TNFb (Tf-in), which efficiently spliced out introns from the neomycin resistance gene (22), and c) the ribozyme self-splicing intron (group I intron) of *Tetrahymena thermophila* (Rb-in) (23). **3)** we selected the best potential sites for intron insertion to increase intron/exon boundaries that are accessible to the spliceosome complex and devoid of interfering secondary structures as previously described (6). Specifically, we inserted the shGb and Tf introns downstream to CAG sequences of the reversed *nluc* gene, and optimized predicted RNA folding by mfold (6). Variants containing intron(s) insertion with the highest numbers of unpaired nucleotides at splice donor (SD), splice acceptor (SA) sites, and the adjacent Nluc sequence are highlighted in green in Figure S1. and **4)** we introduced several nonsense mutations at the 3’-coding region of the reversed *nluc* gene (shown in purple in Figure S1) to reduce the pairing with SD, as well as other nonsense mutations to generate complementary sequence to the internal guide sequence of Rb (both are highlighted in yellow in Figure S1) that are required for efficient ribozyme splicing (24). We applied this design to build 2 types of plasmids: 1) pUCHR-based second-generation lentiviral vector containing the CMV promoter, and 2) pTwist-based third-generation lentiviral vector containing the EF1a-HTLV hybrid promoter (Fig. 1D).

### Transduction and splicing efficiencies of new reporter vectors in 293T cells

We first tested the ability of our vectors to detect cell-cell transmission in 293T cells and in coculture of 293T and Cf2-Th/CD4^+^CCR5^+^ cells in the presence, and as a control, absence of the vesicular stomatitis virus G protein, which mediates efficient entry into a variety of cells (Figure 2A). pUCHR-shGb-inNluc resulted in high cell-cell transmission readout and high background whereas pUCHR-Tf-inNluc exhibited the highest signal/noise ratio. Notably, the newly designed pUCHR-shGb-inNluc was ∼4 orders of magnitude more sensitive than the original inLuc-mR construct but displayed increased Env-independent activity (Figure 2B). Similar results were obtained using HIV-1_AD8_ Envs to mediate cell-cell transmission in a coculture of 293T (producing) cells and Cf2-Th/CD4^+^CCR5^+^ (target) cells (Figure 2C). Low transfection efficiency into Cf2-Th/CD4^+^CCR5^+^ most likely led to dominating cell-cell transmission from 293T to Cf2-Th/CD4^+^CCR5^+^ cells. As expected, HIV-1_AD8_ Envs were less efficient than VSV-G in mediating viral entry and resulted in a decreased signal/noise ratio. Surprisingly, background levels could be significantly decreased by the non-nucleoside reverse transcriptase inhibitor Etravirine, suggesting that the source of background was related to Env-independent cell-cell transmission into target cells rather than background Nluc expression in the producing cells in this coculture. We determined the efficiency of reporter RNA splicing and packaging by measuring the ratio of spliced to total RNA packaged into virions. Splicing efficiencies generally correlated with the cell-cell transmission measurements (Figure 2D).

**Figure 2.**
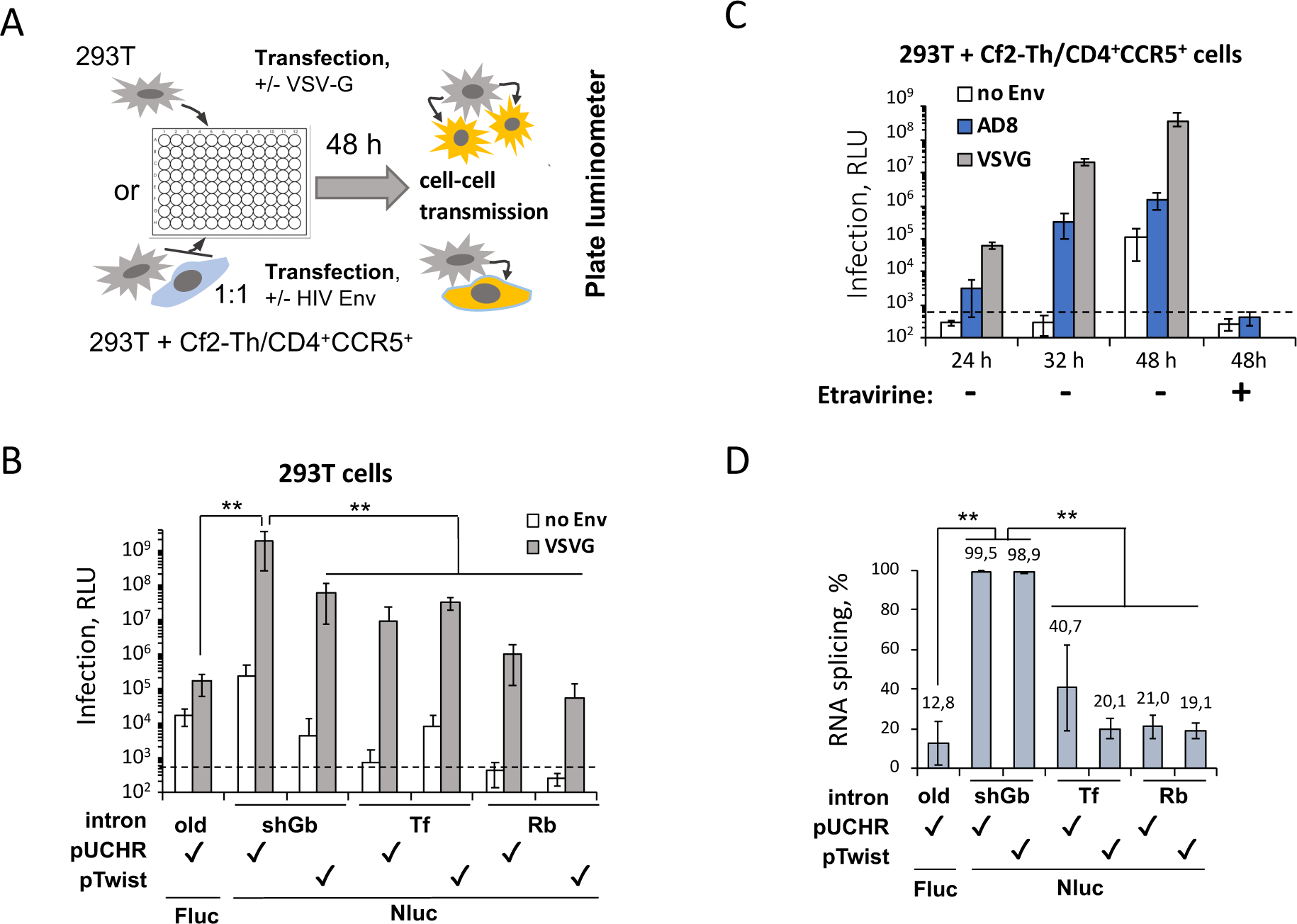
Optimizing the in-Nluc reporter vectors for detection of cell-cell transmission in 293T cells. **(A)** A scheme of assay set up for detection of cell-cell transmission in 293T cells or 293T-Cf2-Th/CD4^+^CCR5^+^ coculture. (**B**) Levels of HIV-1 cell-cell transmission to 293T cells, measured using different vectors and *luc* genes, 48 hours after co-transfecting 293T cells with psPAX2 packaging and indicated reporter plasmids with or without VSV-G expression plasmid. (**C**) Levels of cell-cell transmission from 293T to Cf2-Th/CD4^+^CCR5^+^ cells measured by pUCHR-shGb-inNluc vector at indicated time points post transfection in the presence or absence of indicated Envs and the non-nucleoside reverse transcriptase inhibitor Etravirine. Dashed line indicates background level of Nluc activity. (**D**) Ratio of spliced and unspliced reporter RNAs packaged into virions was quantified by RT-qPCR (see Methods for details). Numbers indicate the percentage spliced RNA. Results (**B**-**D**) are representative of one out of three independent experiments. **, Student’s *t* test *P* value < 0.01. competition of the reporter RNA with the viral genomic RNA (from the full-length HIV-1 plasmids), the reporter RNA was still efficiently packaged into virions. Cell-cell transmission detected by pUCHR-shGb-inNluc exhibited a higher readout among the different inNluc reporter vectors when cotransfected with pNL(AD8) (Figure 3E). And overall readout correlated with the efficiency of the spliced reporter RNA to be packaged into HIV-1 virions (Figure 3F, orange histograms). Notably, while shGb intron splicing is equally efficient when incorporated in pUCHR and pTwist vectors (Figure 2D), the pUCHR provided an about twofold

Overall, shGb intron-based vectors supported high efficiency splicing of Nluc RNA, high sensitivity in a 96-well format (2×10^4^ of 293T transfected with 22 ng of plasmid DNA), and low background in producing cells.

### Quantification of HIV-1 cell-cell transmission and free virus infection in human T cell lines with the new reporter vectors

We next tested the ability of our reporter system to measure cell-cell transmission in T-cells, which more closely reflects the authentic replication of HIV-1 in primary CD4^+^ T cells. We used CEM as producing cells and SupT1.CCR5 as target cells (in a 12-well and 96-well format) to test different combinations of reporter, packaging, and Env-expression plasmids and optimize the system (Figure 3A). Vectors containing the shGb intron cotransfected with Gag-Pol expressing plasmid (psPAX2) and HIV-1 Envs exhibited the highest level of Nluc expression, but, in contrast to 293T cells, pTwist-based plasmid was more sensitive than pUCHR-based plasmid (Figure 3B). We detected only minimal background in the T-cell coculture system. Interestingly, the cotransfection of the pUCHR-shGb-inNluc reporter with a full-length HIV-1 vector pNL41′Env containing only a frame-shift mutation in *env* gene led to a higher level of cell-to-cell transmission in T cells compared to two versions of packaging plasmids (pCMV-dR8.2 and psPAX2) and even higher readout was measured when we used a plasmid containing a full-length molecular clone pNL4(AD8) (Figure 3D). Thus, despite potential higher level of total RNA incorporation into virions compared to analogous pTwist vector, and more efficient incorporation led to higher signal. Consistent with previous reports, free virus infection of HIV-1_AD8_ was ∼15-fold less efficient (Figure 3G) than cell-cell transmission.

**Figure 3.**
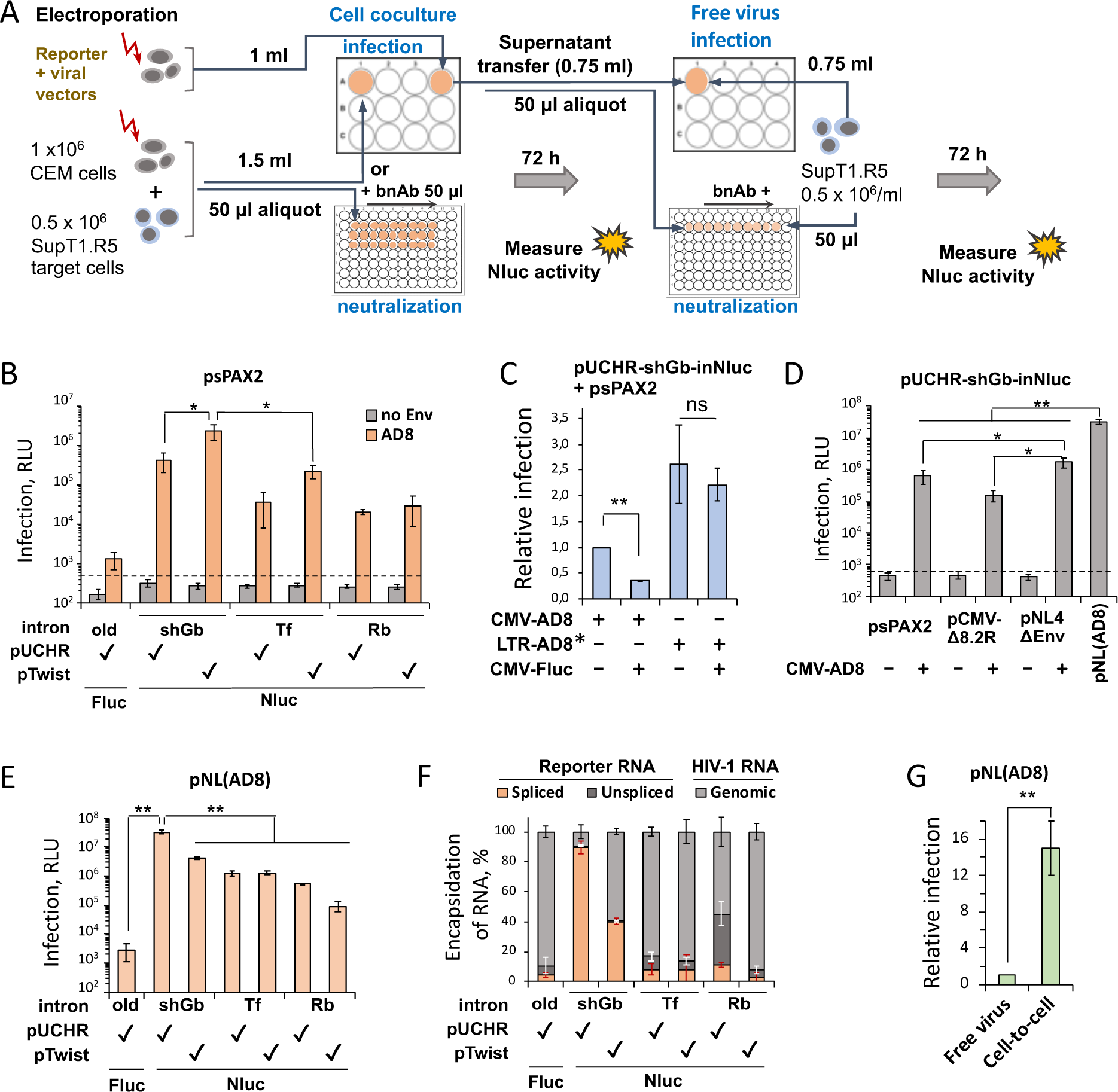
Cell-cell transmission in CD4^+^ T cell lines detected by the inNluc reporter vectors. (**A**) A workflow for quantification of cell-cell transmission from CEM to SupT1.CCR5 cells by the inNluc reporter vectors and, in parallel, of free virus infection under identical conditions. (**B**) Cell-cell transmission measured by different reporter vectors cotransfected with the packaging (psPAX2) and HIV-1_AD8_ Env-expressing plasmids, in which transgenes are transcribed from the CMV promoter. (**C**) Effects of CMV-driven transgene expression on the cell-cell transmission readout. Cells were co-transfected with pUCHR-CMV-shGb-inNluc reporter vector, packaging plasmid psPAXS2 and different HIV-1_AD8_ Env-expressing plasmids with or without Fluc-expressing plasmid pGL3-CMV. *, 3’ end of the *env* gene in LTR-AD8 is driven from NL4-3 *env*. (**D**) Effects of different packaging vectors or full-length HIV-1 with deleted *env* on the cell-cell transmission readout. (**E**) Measurements of cell-cell transmission mediated by the molecular clone pNL4(AD8) using the indicated reporter vectors. (**F**) Ratio of spliced reporter RNA, unspliced reporter RNA, and viral genomic RNA packaged into virions produced in 293T cells after cotransfection with pNL(AD8) and the indicated reporters. (G) In parallel comparison of HIV-1_AD8_ cell-cell transmission and free virus infectivity. Half of the supernatant from cell-cell transmission assay containing free virus was cleared and used to infect SupT1.R5 target cells. *, *P* value < 0.05; **, *P* value < 0.01; both *P* values were calculated by two-tailed Student’s t-test.

### Detection of HIV-1 cell-cell transmission between primary human CD4^+^ T cells

We next selected the two most efficient reporter vectors (containing the shGb intron in pUCHR and pTwist backbones) and tested their ability to measure cell-cell transmission in primary CD4+ T cells. For these experiments, we also generated a pUCHR-based vector in which we replaced the CMV promoter by EF1a-HTLV-1 hybrid promotor to study the effect of promoter on the measurements in primary cells. We isolated CD4^+^ T cells from 2 donors, activated the cells and electroporated as previously described (25). All subsequent steps were performed as described for the T cell lines but transfected cells were cocultured with autologous primary CD4^+^ T cells instead of the heterologous coculture using SupT1.CCR5 cells. pUCHR-shGb-inNluc vectors containing either CMV or EF1a promoter, cotransfected with pNL4-3 (a plasmid that contains the NL4-3 molecular clone; X4 tropic), resulted in high and comparable readout of cell-cell transmission (Figure 4A). In comparison, the pTwist-based vector demonstrated ∼ 6.7-fold lower level of Nluc activity in primary CD4^+^ T cells, consistent with the low efficient packaging of RNA transcribed from the pTwist reporter into HIV-1 virions (Figure 3F). We also detected comparable cell-cell transmission of pNL4-3 and an R5-tropic molecular clone NL(AD8) despite typically lower levels of CCR5 than CXCR4 expected on activated CD4^+^ T cells. Single-round HIV-1 vectors could support sensitive detection of cell-cell transmission in primary CD4^+^ T cells as well, using the reporter plasmid together with packaging plasmid and the dual-tropic HIV-1_KB9_ Envs. Our system exhibited minimal background without HIV-1 Envs for all packaging plasmids (Figure 4B) and we detected a differential readout of cell-cell transmission using different packaging plasmids that ranked as psPAX2< pCMV-d8.2R< pNL4-3. Notably, unlike the results with T cell lines, the pCMV-d8.2R packaging vector in primary cells provided a higher level of HIV-1 infection than the psPAX2 vector. This may reflect the requirement of accessory genes for efficient HIV replication in primary cells which are deleted in the psPAX2 vector.

**Figure 4.**
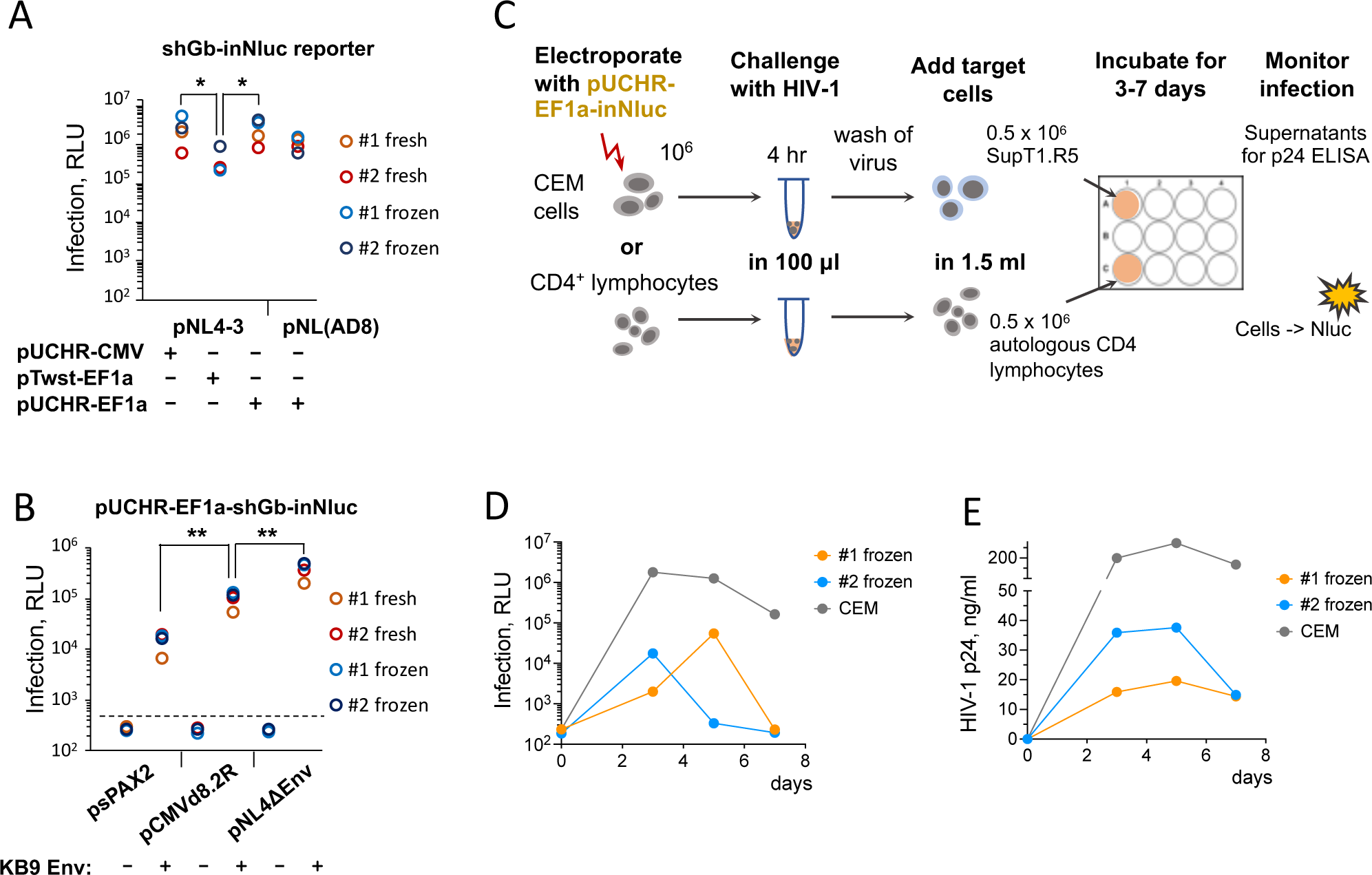
Detection of HIV-1 cell-cell transmission in human primary CD4+ T cells by different reporter vectors. (**A-B**) Cell-cell transmission in primary CD4+ T cells after co-transfection of different reporter vectors containing the shGB-inNluc gene together with plasmids containing full-length HIV-1 provirus (**A**) or with plasmids containing HIV-1 packaging genes (**B**). Data in (**A**) and (**B**) were obtained using fresh or frozen/thawed PBMCs from 2 donors (#1 and #2). (**C**) A scheme of experimental set up for monitoring cell-cell transmission during HIV-1 replication. **D-E,** Kinetics of cell-cell transmission as detected by Nluc expression during HIV-1 replication (**D**) and kinetics of HIV-1 replication in the same cells detected by p24 release into the supernatant (**E**). CEM or primary CD4+ T cells were transfected with pUCHR-EF1a-shGb-inNluc reporter vector and then infected with 300 ng HIV-1_NL4-3_. *, *P* value < 0.05; **, *P* value < 0.01; both *P* values were calculated by two-tailed Student’s t-test.

To study HIV-1 cell-cell transmission after virus inoculation, CEM cells or primary CD4^+^ T cells were electroporated with the reporter vector pUCHR-EF1a-shGb-inNluc only. Transfected cells were then incubated with the concentrated HIV-1, washed, and mixed with target cells, SupT1.CCR5 or primary CD4+ T cells, respectively (Figure 4C). Our reporter vector could detect cell-cell transmission in cell lines and primary CD4+ T cells within 3-5 days after inoculation of 300 ng of HIV-1_NL4-3_ although detection was one to two orders of magnitude lower compared to co-transfection of all viral vectors. Nluc expression in target cells correlated with p24 in the supernatants of infected cells (Figure 4E).

In conclusion, the high sensitivity of the new in-Nluc vectors provides robust detection of HIV-1 cell-cell transmission in activated primary CD4^+^ T cell cocultures by co-transfecting with HIV-1 molecular clones, single-round replication vectors, or by inoculating HIV-1 virions.

### Sensitivity of cell-cell transmission to broadly neutralizing antibodies (bnAbs) against HIV-1 Envs

High sensitivity and low background levels detected with the pUCHR-EF1a-shGb-inNluc reporter vector in T cells provided a wide dynamic range for accurate quantification of cell-cell transmission sensitivity to neutralization by bnAbs. We measured the sensitivity to bnAbs against different HIV-1 Env vulnerability sites (CD4 binding site, gp120 V3 glycan, and MPER) and compared the inhibition efficiency of HIV-1 transmission (expressed as IC_50_; half-maximal inhibitory concentration) to the inhibition efficiency of free viruses present in the supernatant of these cells. To produce free viruses under identical conditions, transfected CEM cells without target cells were incubated in parallel to the CEM-SupT1.CCR5 coculture for 3 days and then virion-containing supernatant was added to diluted bnAbs and SupT1.CCR5 target cells (a single replicate) and the plate was incubated for 3 days. Consistent with several previous reports bnAbs were significantly less efficient in blocking cell-cell transmission than neutralization of free virions in supernatant (26); overall we measured at least one order of magnitude difference in neutralization efficiency for all tested bnAbs (Figure 5 A and C). HIV-1 cell-cell transmission was most resistant to VRC01 (IC_50_= 20 μl/ml) while free viruses were still sensitive to VRC01 with an IC_50_=0.12 μl/ml, which was >2 orders of magnitude more efficient than blocking transmission between cells. We also quantify the sensitivity of HIV-1 cell-to-cell transmission in primary CD4^+^ T cells to bnAbs (Figure 5B) and detected the same general pattern of relative resistance of cell-cell transmission to bnAb inhibition (Fig. 5 C and Suppl Table S3 for readout levels).

**Figure 5.**
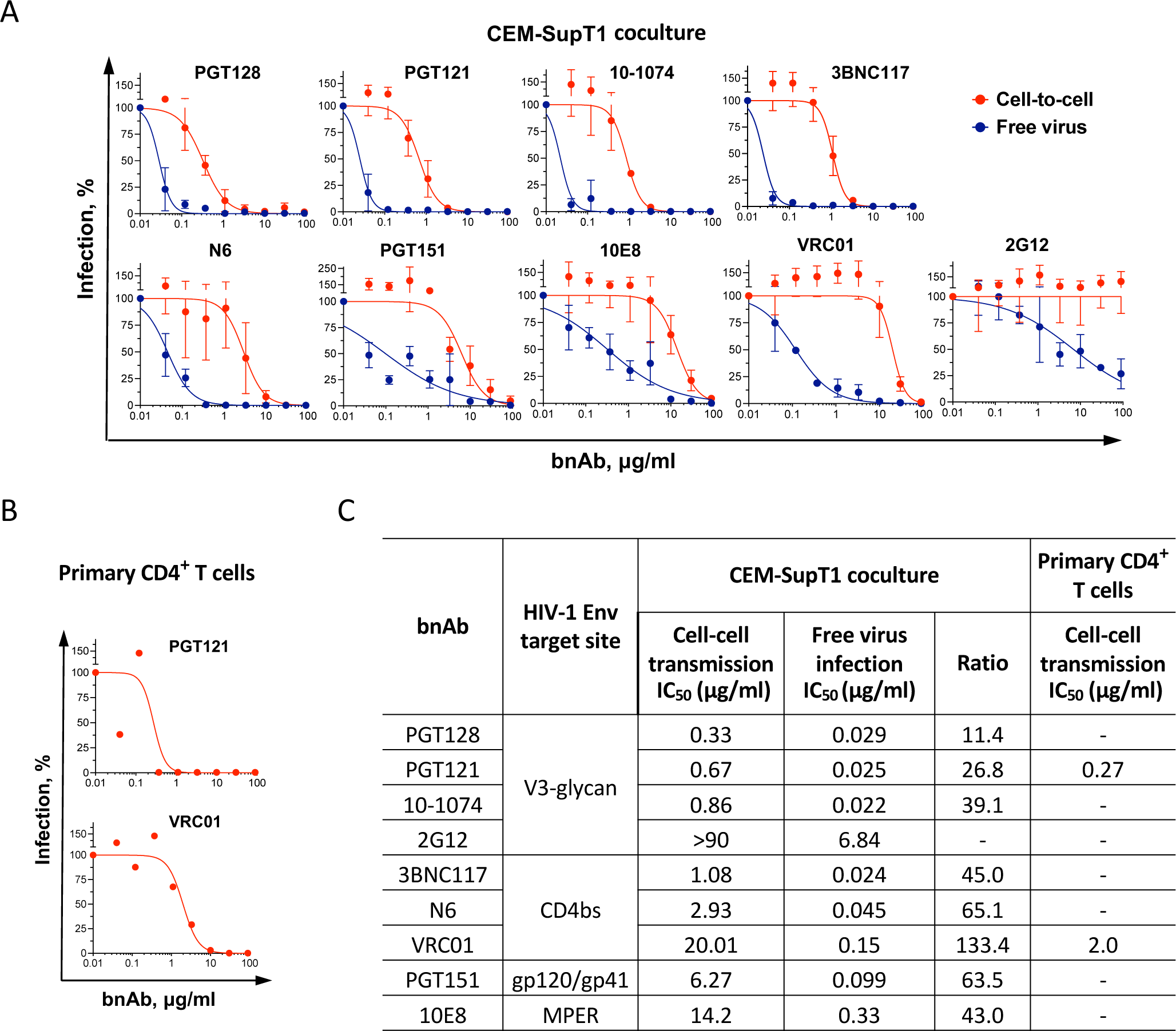
Sensitivity of HIV-1 cell-cell transmission in CD4^+^ T cells to bnAbs. (**A**) Dose-response curves of HIV-1 cell-to-cell transmission and free virus infection in CEM-SupT1 cell system in the presence of 7 individual bnAbs targeting different sites of AD8 Envs. CEM cells were co-transfected with pUCHR-EF1a-shGb-inNluc, pNL4ΔEnv, and LTR-AD8 plasmid DNA and used in neutralization of two modes of HIV-1 transmission as described in Method section. Assays on T cell lines were performed twice in duplicates and data are presented as average values with the standard deviations. (**B**) Sensitivity of cell-to-cell transmission between primary CD4^+^ T cells to the indicated bnAbs. Activated CD4+ lymphocytes were electroporated with the same reporter vector and pNL(AD8) plasmid. A mixture of transfected and non-transfected autologous CD4^+^ T cells was added to the diluted bnAbs in one replica. PGT 121 sensitivity of HIV-1_AD8_ transmission in primary CD4^+^ T cells from donor #2 and VRC01 sensitivity of HIV-1_AD8_ transmission in primary CD4^+^ T cells from donor #1 are shown (from a single experiment). (**C**) Calculated IC_50_ values for HIV-1_AD8_ sensitivity to each bnAbs tested with free virus or during cell-to-cell transmission.

Thus, our new reporter vector can robustly measure sensitivity of HIV-1 cell-to-cell transmission to bnAbs for both CD4^+^ T cell lines and primary CD4^+^ T cells.

## DISCUSSION

Quantitative measurement of HIV cell-to-cell transmission using reporter vectors is challenging. Reversed and intron-inactivated reporter genes (e.g. *fluc*) allow distinguishing between transfected and newly infected cells in cell cocultures. However, currently available reporter vectors that use the reverse firefly luciferase gene and contain an intron have substantial drawbacks due to packaging of HIV-1 genome containing unspliced reporter gene (5,6). These vectors allow measurements of HIV-1 cell-cell transmission between lymphoid cell lines, as well as in cocultures of Raji cells and PBMCs, but detection of transmission between primary CD4+ T cells is still very limited (6). Alternative in-*Gluc* reporter vectors, which were initially designed for detection of cell-cell transmission of murine leukemia virus (MLV) (27) and subsequently adapted for HIV-1 (28), are based on secreted *Gaussia* luciferase. These vectors exhibit three orders of magnitude higher signal compared to the firefly luciferase (29) but high signal is associated with elevated background levels due to oxidation of the coelenterazine substrate by components in FBS typically present in culture media (30). As a result, the signal- to-background ratio using these vectors is reduced and they have been mainly used to measure cell-cell transmission in highly transfectable 293T cells to highly permissive cell lines 293T.CD4.CXCR4, HeLa TZMblue, MT4 (28) or A3.01.CCR5 (26).

Here, we addressed current limitations on cell-cell transmission reporter vectors by designing and building ultra-sensitive assay to measure HIV-1 cell-to-cell transmission. We optimized the length of shRNA-containing intron without compromising splicing efficiency; and we codon optimized sequences that are predicted to involve in RNA secondary structure at splice acceptor (SA) and splice donor (SD) sites to prevent potential interference. This approach led to very efficient *nluc* gene splicing (Figure 2E), which together with superior brightness and minimal background, provide unprecedent high sensitivity and wide dynamic range of the NLuc reporter vectors. As a result, in CEM-SupT1 coculture, co-transfection of pUCHR-shGb-inNluc with pNL4(AD8) generated a signal/noise ratio of >80,000 (Figure 3D). High sensitivity and wide dynamic range allowed us to robustly measure the sensitivity of HIV-1 cell-cell transmission to neutralizing antibodies using CD4+ T cells in 96-well plate format and compare this activity side-by-side with HIV-1 sensitivity to neutralizing antibodies in parallel using the same cell system (Figure 5).

Our improved and highly sensitive technology can immediately be applied to answer important research questions. As our reporter vector can detect HIV-1 replication in cocultures of primary CD4^+^ T cells (Figure 4A and B), this system can be utilized to measure the ability of different HIV-1 strains from people living with HIV-1 (PLWH) to transmit between their primary CD4^+^ T cells ex vivo. Current anti-retroviral therapy decreases HIV-1 viral load to undetectable levels in most treated PLWH and therefore isolating CD4+ T cells from PLWH and transfecting them with the NLuc reporter vector with subsequent activation of HIV-1 replication will reveal cell-cell transmission efficiency of different HIV-1 Envs.

More importantly, our new ultrasensitive tool can accurately measure the sensitivity of cell-cell transmission to antibody neutralization. Due to broad coverage and high potency, bnAb have been or are now being tested as therapeutic and prevention modalities in several completed and ongoing clinical trials. However, in some cases, despite high level of bnAbs (i.e., VRC01 or VRC07) in the blood of treated patients, HIV-1 is rebounding but the recrudescing strains are still sensitive *in vitro* to VRC01 or VRC07 and they do not exhibit any evidence of viral evolution to resist bnAbs (31,32). One potential mechanism of HIV-1 resistance may be increased ability to transmit between cells *in vivo* (33), a process which is relatively resistant to most bnAbs and particularly resistant to VRC01. Thus, we are now in the process of obtaining resistant HIV-1 strains from immunotherapy clinical trials to study their ability to transmit between primary CD4^+^ T cells and to investigate the relationship between HIV-1 resistance to bnAbs and cell-cell transmission efficiency *ex vivo*.

In addition, application of the reporter vectors is specific but not limited to HIV-1. They can be applied for evaluation of humoral response against other viruses, specifically, SARS-CoV-2 whose cell-to-cell transmission has been reported to be resistant to neutralization (34,35). The EF1a-shGb-inNluc-pA expression cassette can be subcloned into the other retroviral vectors, such as vectors supporting HTLV-I and MLV, to study cell-to-cell transmission at a new level of sensitivity. The algorithm for shGb intron insertion described here can be applied to create different selection genes with a high level of reconstitution, to detect, for instance, retro-transposition events.

Overall, our improved reporter will provide new opportunities to study the mechanisms of viral cell-cell transmission, to categorize different HIV-1 strains according to their cell-cell transmission, and to investigate the cell-cell transmission sensitivity to antibody neutralization. These directions studied in primary cells are expected to provide new insights into relevant and complex biological processes.

## AUTHOR CONTRIBUTIONS

D.M. and A.H. conceived and designed the study. D.M. performed the experiments and analyzed the data. D.M. wrote the first draft of the manuscript and A.H. edited and finalized the manuscript.

## ACKNOWLEDGEMENTS

We thank Durgadevi Parthasarathy (University of Minnesota) for maintaining cells and providing plasmids; Branden Moriarity and Evan Kleinboehl (University of Minnesota) for providing access to the Neon electroporation system; and James Hoxie (University of Pennsylvania) for the SupT1.CCR5 cells. We also thank the NIH AIDS Reagent program for providing the following antibodies: VRC01, N6, 2G12, PGT121, PGT128, 10-1074, and 10E8; and the plasmids pNL4-3 and pNL(AD8).

## DATA AVAILABILITY

Data is available upon request.

## SUPPLEMENTARY DATA

Supplementary Data are available at NAR online.

---

**Table S1.**
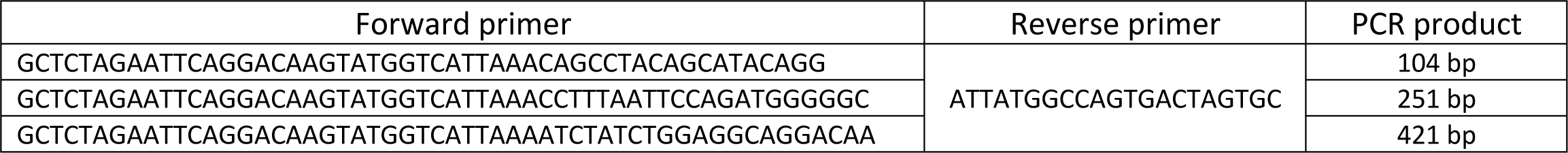
Primers used to shorten the human ψ-intron.

**Table S2.**
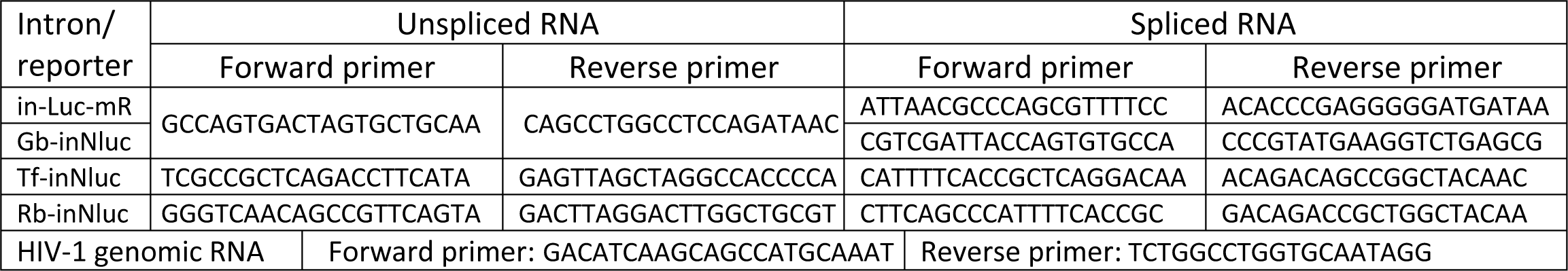
RT-qPCR primers for quantification of spliced/unspliced reporter and HIV-1 genomic RNAs in viral particles.

**Table S3.** Average RLU values obtained in a 96-well plate HIV-1 cell-to-cell neutralization assay.

| bnAb | CEM-SupT1 cell system |  |  | Primary CD4+ T cells |  |  |
| --- | --- | --- | --- | --- | --- | --- |
|  | 0 µg/ml | 1.1 µg/ml | 30 µg/ml | 0 µg/ml | 1.1 µg/ml | 30 µg/ml |
| PGT128 | 15,294,565 | 1,640,440 | 871,231 |  |  |  |
| PGT121 | 16,582,305 | 5,863,471 | 9,948 | 268,766 | 1,133 | 587 |
| 10-1074 | 14,807,042 | 4,730,994 | 1,613 |  |  |  |
| 3BNC117 | 13,035,579 | 6,000,224 | 1,310 |  |  |  |
| N6 | 16,582,305 | 14,350,783 | 57,669 |  |  |  |
| PGT151 | 12,908,383 | 14,467,548 | 1,661,023 |  |  |  |
| 10E8 | 14,405,412 | 13,157,143 | 4,261,206 |  |  |  |
| VRC01 | 6,631,937 | 9,297,742 | 1,186,488 | 231,985 | 181,790 | 1,042 |
| 2G12 | 12,377,928 | 18,545,674 | 13,374,130 |  |  |  |

**Figure S1.**
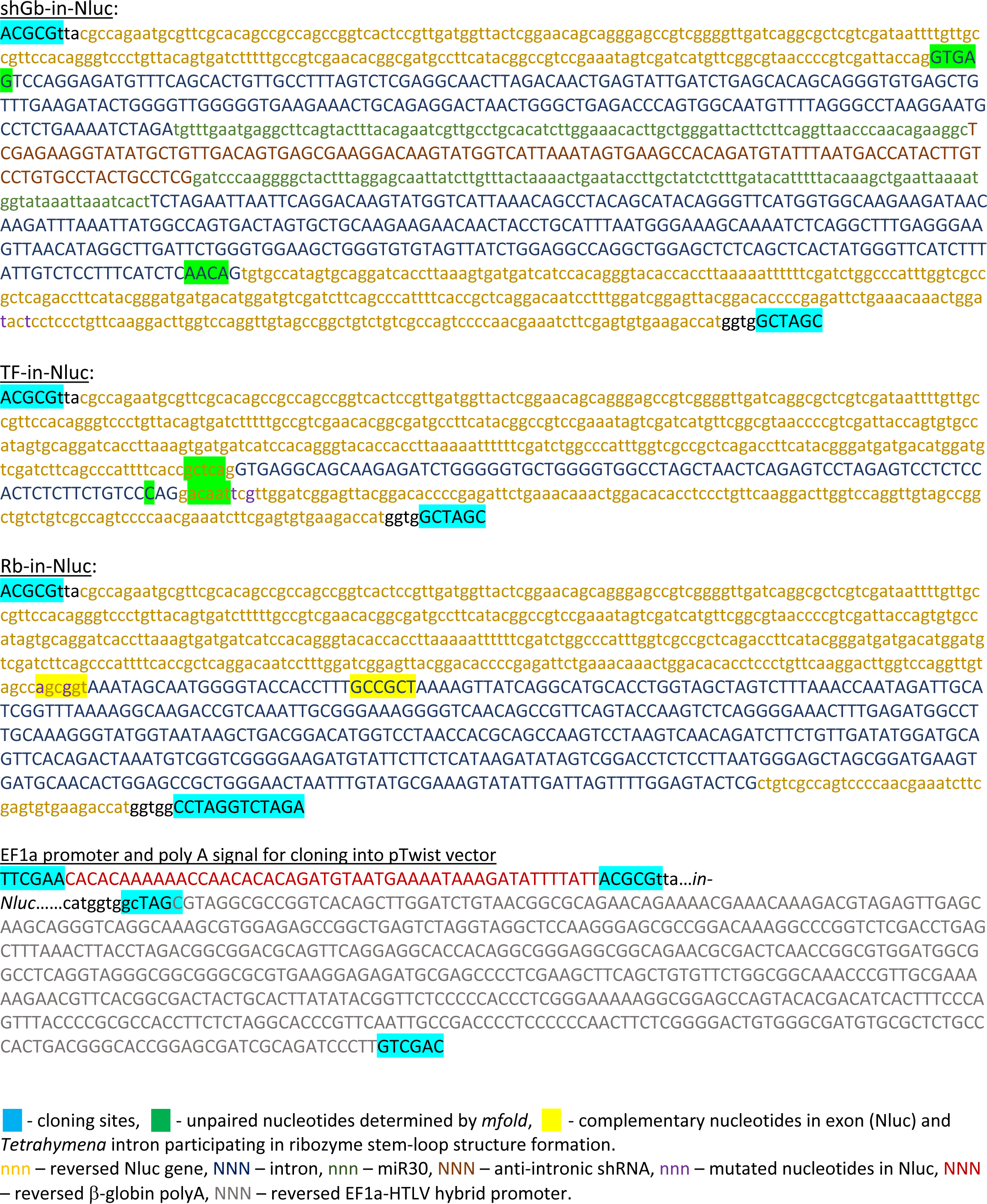
Sequences used to construct HIV-1 replication-dependent vectors with nanoluciferase gene.

## REFERENCES

1. Jolly, C., Kashefi, K., Hollinshead, M. and Sattentau, Q.J. (2004) HIV-1 cell to cell transfer across an Env-induced, actin-dependent synapse. J Exp Med, 199, 283–293.

2. Malbec, M., Porrot, F., Rua, R., Horwitz, J., Klein, F., Halper-Stromberg, A., Scheid, J.F., Eden, C., Mouquet, H., Nussenzweig, M.C. et al. (2013) Broadly neutralizing antibodies that inhibit HIV-1 cell to cell transmission. J Exp Med, 210, 2813–2821.

3. Chen, P., Chen, B.K., Mosoian, A., Hays, T., Ross, M.J., Klotman, P.E. and Klotman, M.E. (2011) Virological synapses allow HIV-1 uptake and gene expression in renal tubular epithelial cells. J Am Soc Nephrol, 22, 496–507.

4. Chen, P., Hübner, W., Spinelli, M.A. and Chen, B.K. (2007) Predominant mode of human immunodeficiency virus transfer between T cells is mediated by sustained Env-dependent neutralization-resistant virological synapses. J Virol, 81, 12582–12595.

5. Mazurov, D., Ilinskaya, A., Heidecker, G., Lloyd, P. and Derse, D. (2010) Quantitative comparison of HTLV-1 and HIV-1 cell-to-cell infection with new replication dependent vectors. PLoS Pathog, 6, e1000788.

6. Shunaeva, A., Potashnikova, D., Pichugin, A., Mishina, A., Filatov, A., Nikolaitchik, O., Hu, W.S. and Mazurov, D. (2015) Improvement of HIV-1 and Human T Cell Lymphotropic Virus Type 1 Replication-Dependent Vectors via Optimization of Reporter Gene Reconstitution and Modification with Intronic Short Hairpin RNA. J Virol, 89, 10591–10601.

7. Wu, X., Yang, Z.Y., Li, Y., Hogerkorp, C.M., Schief, W.R., Seaman, M.S., Zhou, T., Schmidt, S.D., Wu, L., Xu, L. et al. (2010) Rational design of envelope identifies broadly neutralizing human monoclonal antibodies to HIV-1. Science, 329, 856–861.

8. Ofek, G., Guenaga, F.J., Schief, W.R., Skinner, J., Baker, D., Wyatt, R. and Kwong, P.D. (2010) Elicitation of structure-specific antibodies by epitope scaffolds. Proc Natl Acad Sci U S A, 107, 17880–17887.

9. Walker, L.M., Phogat, S.K., Chan-Hui, P.Y., Wagner, D., Phung, P., Goss, J.L., Wrin, T., Simek, M.D., Fling, S., Mitcham, J.L. et al. (2009) Broad and potent neutralizing antibodies from an African donor reveal a new HIV-1 vaccine target. Science, 326, 285–289.

10. Huang, J., Ofek, G., Laub, L., Louder, M.K., Doria-Rose, N.A., Longo, N.S., Imamichi, H., Bailer, R.T., Chakrabarti, B., Sharma, S.K. et al. (2012) Broad and potent neutralization of HIV-1 by a gp41-specific human antibody. Nature, 491, 406–412.

11. Huang, J., Kang, B.H., Ishida, E., Zhou, T., Griesman, T., Sheng, Z., Wu, F., Doria-Rose, N.A., Zhang, B., McKee, K. et al. (2016) Identification of a CD4-Binding-Site Antibody to HIV that Evolved Near-Pan Neutralization Breadth. Immunity, 45, 1108–1121.

12. Zhou, T., Georgiev, I., Wu, X., Yang, Z.Y., Dai, K., Finzi, A., Kwon, Y.D., Scheid, J.F., Shi, W., Xu, L. et al. (2010) Structural basis for broad and potent neutralization of HIV-1 by antibody VRC01. Science, 329, 811–817.

13. Flemming, J., Wiesen, L. and Herschhorn, A. (2018) Conformation-Dependent Interactions Between HIV-1 Envelope Glycoproteins and Broadly Neutralizing Antibodies. AIDS Res Hum Retroviruses, 34, 794–803.

14. Herschhorn, A., Gu, C., Espy, N., Richard, J., Finzi, A. and Sodroski, J.G. (2014) A broad HIV-1 inhibitor blocks envelope glycoprotein transitions critical for entry. Nat Chem Biol, 10, 845–852.

15. Herschhorn, A., Gu, C., Moraca, F., Ma, X., Farrell, M., Smith, A.B., 3rd, Pancera, M., Kwong, P.D., Schön, A., Freire, E. et al. (2017) The β20-β21 of gp120 is a regulatory switch for HIV-1 Env conformational transitions. Nat Commun, 8, 1049.

16. Ratnapriya, S., Chov, A. and Herschhorn, A. (2020) A Protocol for Studying HIV-1 Envelope Glycoprotein Function. STAR Protoc, 1, 100133.

17. Harris, M., Ratnapriya, S., Chov, A., Cervera, H., Block, A., Gu, C., Talledge, N., Mansky, L.M., Sodroski, J. and Herschhorn, A. (2020) Slow Receptor Binding of the Noncytopathic HIV-2(UC1) Envs Is Balanced by Long-Lived Activation State and Efficient Fusion Activity. Cell Rep, 31, 107749.

18. Herschhorn, A., Ma, X., Gu, C., Ventura, J.D., Castillo-Menendez, L., Melillo, B., Terry, D.S., Smith, A.B., 3rd, Blanchard, S.C., Munro, J.B. et al. (2016) Release of gp120 Restraints Leads to an Entry-Competent Intermediate State of the HIV-1 Envelope Glycoproteins. mBio, 7.

19. Ratnapriya, S., Braun, A.R., Cervera Benet, H., Carlson, D., Ding, S., Paulson, C.N., Mishra, N., Sachs, J.N., Aldrich, C.C., Finzi, A. et al. (2022) Broad Tricyclic Ring Inhibitors Block SARS-CoV-2 Spike Function Required for Viral Entry. ACS Infect Dis, 8, 2045–2058.

20. England, C.G., Ehlerding, E.B. and Cai, W. (2016) NanoLuc: A Small Luciferase Is Brightening Up the Field of Bioluminescence. Bioconjug Chem, 27, 1175–1187.

21. Loh, J.M. and Proft, T. (2014) Comparison of firefly luciferase and NanoLuc luciferase for biophotonic labeling of group A Streptococcus. Biotechnol Lett, 36, 829–834.

22. Esnault, C., Casella, J.F. and Heidmann, T. (2002) A Tetrahymena thermophila ribozyme-based indicator gene to detect transposition of marked retroelements in mammalian cells. Nucleic Acids Res, 30, e49.

23. Hasegawa, S., Jackson, W.C., Tsien, R.Y. and Rao, J. (2003) Imaging Tetrahymena ribozyme splicing activity in single live mammalian cells. Proc Natl Acad Sci U S A, 100, 14892–14896.

24. Zufferey, R., Dull, T., Mandel, R.J., Bukovsky, A., Quiroz, D., Naldini, L. and Trono, D. (1998) Self-inactivating lentivirus vector for safe and efficient in vivo gene delivery. J Virol, 72, 9873–9880.

25. Maslennikova, A., Kruglova, N., Kalinichenko, S., Komkov, D., Shepelev, M., Golubev, D., Siniavin, A., Vzorov, A., Filatov, A. and Mazurov, D. (2022) Engineering T-Cell Resistance to HIV-1 Infection via Knock-In of Peptides from the Heptad Repeat 2 Domain of gp41. mBio, 13, e0358921.

26. Reh, L., Magnus, C., Schanz, M., Weber, J., Uhr, T., Rusert, P. and Trkola, A. (2015) Capacity of Broadly Neutralizing Antibodies to Inhibit HIV-1 Cell-Cell Transmission Is Strain- and Epitope-Dependent. PLoS Pathog, 11, e1004966.

27. Aloia, A.L., Duffy, L., Pak, V., Lee, K.E., Sanchez-Martinez, S., Derse, D., Heidecker, G., Cornetta, K. and Rein, A. (2013) A reporter system for replication-competent gammaretroviruses: the inGluc-MLV-DERSE assay. Gene Ther, 20, 169–176.

28. Zhong, P., Agosto, L.M., Ilinskaya, A., Dorjbal, B., Truong, R., Derse, D., Uchil, P.D., Heidecker, G. and Mothes, W. (2013) Cell-to-cell transmission can overcome multiple donor and target cell barriers imposed on cell-free HIV. PLoS One, 8, e53138.

29. Tannous, B.A., Kim, D.E., Fernandez, J.L., Weissleder, R. and Breakefield, X.O. (2005) Codon-optimized Gaussia luciferase cDNA for mammalian gene expression in culture and in vivo. Mol Ther, 11, 435–443.

30. Zhao, H., Doyle, T.C., Wong, R.J., Cao, Y., Stevenson, D.K., Piwnica-Worms, D. and Contag, C.H. (2004) Characterization of coelenterazine analogs for measurements of Renilla luciferase activity in live cells and living animals. Mol Imaging, 3, 43–54.

31. Cale, E.M., Bai, H., Bose, M., Messina, M.A., Colby, D.J., Sanders-Buell, E., Dearlove, B., Li, Y., Engeman, E., Silas, D. et al. (2020) Neutralizing antibody VRC01 failed to select for HIV-1 mutations upon viral rebound. J Clin Invest, 130, 3299–3304.

32. Bar, K.J., Sneller, M.C., Harrison, L.J., Justement, J.S., Overton, E.T., Petrone, M.E., Salantes, D.B., Seamon, C.A., Scheinfeld, B., Kwan, R.W. et al. (2016) Effect of HIV Antibody VRC01 on Viral Rebound after Treatment Interruption. N Engl J Med, 375, 2037–2050.

33. Herschhorn, A. (2023) Indirect Mechanisms of HIV-1 Evasion from Broadly Neutralizing Antibodies In Vivo. ACS Infect Dis, 9, 5–8.

34. Kruglova, N., Siniavin, A., Gushchin, V. and Mazurov, D. (2021) Different Neutralization Sensitivity of SARS-CoV-2 Cell-to-Cell and Cell-Free Modes of Infection to Convalescent Sera. Viruses, 13.

35. Zeng, C., Evans, J.P., King, T., Zheng, Y.M., Oltz, E.M., Whelan, S.P.J., Saif, L.J., Peeples, M.E. and Liu, S.L. (2022) SARS-CoV-2 spreads through cell-to-cell transmission. Proc Natl Acad Sci U S A, 119.

